# Identification that ADAM17 mediates proteolytic maturation of calcium channel auxiliary α_2_δ subunits, and enables calcium current enhancement

**DOI:** 10.1101/2021.04.29.441911

**Authors:** Ivan Kadurin, Shehrazade Dahimene, Karen M Page, Joseph I. J. Ellaway, Kanchan Chaggar, Linda Troeberg, Hideaki Nagase, Annette C. Dolphin

## Abstract

The auxiliary α_2_δ subunits of voltage-gated calcium (Ca_V_) channels are key to augmenting expression and function of Ca_V_1 and Ca_V_2 channels, and are also important drug targets in several therapeutic areas, including neuropathic pain. The α_2_δ proteins are translated as pre-proteins encoding both α_2_ and δ, and post-translationally proteolysed into α_2_ and δ subunits, which remain associated as a complex. In this study we have identified ADAM17 as a key protease involved in proteolytic processing of pro-α_2_δ-1 and α_2_δ-3 subunits. We provide three lines of evidence: firstly, proteolytic cleavage is inhibited by chemical inhibitors of particular metalloproteases, including ADAM17. Secondly, proteolytic cleavage of both α_2_δ-1 and α_2_δ-3 is markedly reduced in cell lines by knockout of *ADAM17* but not *ADAM10*. Thirdly, proteolytic cleavage is reduced by the N-terminal active domain of TIMP-3 (N-TIMP-3), which selectively inhibits ADAM17. We have found previously that proteolytic cleavage into mature α_2_δ is essential for the enhancement of Ca_V_ function, and in agreement, knockout of ADAM17 inhibited the ability of α_2_δ-1 to enhance both Ca_V_2.2 and Ca_V_1.2 calcium currents. Thus, our study identifies ADAM17 as a key protease required for proteolytic maturation of α_2_δ-1 and α_2_δ-3, and thus a potential drug target in neuropathic pain.

## INTRODUCTION

Voltage-gated calcium (Ca_V_) channels are essential for multiple physiological functions including neurotransmitter release and muscle contraction, and are also important drug targets in several therapeutic areas, including chronic pain (Catterall, 2000; Zamponi et al., 2015). There are three subtypes of Ca_V_ channel pore-forming α_1_ subunit (Ca_V_1, 2 and 3), of which Ca_V_1 and 2 are associated with auxiliary β and α_2_δ subunits (Flockerzi et al., 1986; Liu et al., 1996; Takahashi et al., 1987; Witcher et al., 1993), which are both important for their function (for review see Dolphin, 2012).

The α_2_δ subunits are extracellular proteins that undergo complex post translational modifications (Figure 1A). A single gene encodes each α_2_δ pre-protein, which is then subject to several processing steps, including glycosyl-phosphatidylinositol (GPI)-anchoring (Davies et al., 2010), extensive glycosylation and proteolytic processing into disulfide-linked α_2_ and δ (Ellis et al., 1988; Jay et al., 1991). The cryo-electron microscopic structure of the skeletal muscle Ca_V_1.1 complex (Wu et al., 2016) shows interaction of α_2_δ-1 with several extracellular loops of the α1 subunit, including a key residue in the first extracellular loop of Domain I, which interacts with the von Willebrand factor (VWA) domain of α_2_δ-1.

**Figure 1:**
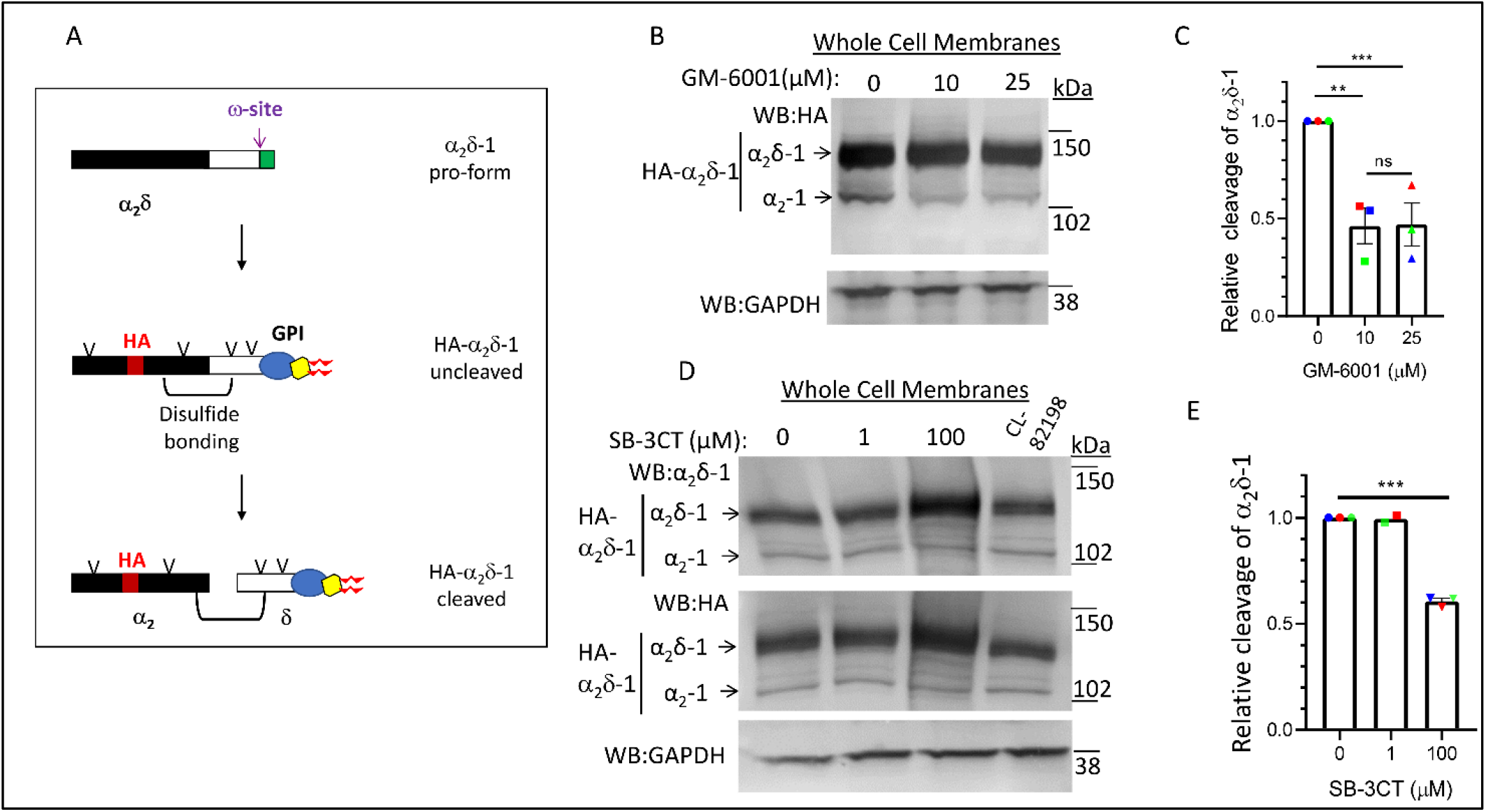
Effect of chemical inhibitors of ADAMs and MMPs on α_2_δ-1 proteolytic cleavage. (A) Diagram of post-translational processing of α_2_δ proteins, including glycosylation (V), GPI anchoring and proteolytic cleavage. It also shows the approximate position of inserted HA tag (red) and disulfide bonding between α_2_ (black) and δ (white). (B) Effect of GM-6001 (0, 10 and 25 µM; lanes 1 – 3, respectively) on cleavage in whole cell membranes of HA-α_2_δ-1 expressed in tsA-201 cells (upper panel: HA immunoblot), de-glycosylated with PNGase-F to allow resolution between pro-α_2_δ-1 (upper band) and the cleaved form, α_2_-1 (lower band). The absolute % cleavage was 12.8 ± 2.0 % in control conditions. Lower panel, loading control: endogenous GAPDH. (C) Quantification of the effect of 10 µM (squares) and 25 µM (triangles) GM-6001 on relative cleavage of α_2_δ-1 (normalized to that under control conditions (circles)). Data are mean ± SEM and individual data in 3 separate experiments, including that in (B), denoted by red, green and blue symbols. Statistical differences determined using 1-way ANOVA and Tukey post hoc test; ** *P* = 0.0084; *** *P* = 0.0090. (D) Effect of SB-3CT (0, 1 and 100 µM; lanes 1 – 3, respectively) and CL-82198 (60 µM; lane 4) on cleavage in whole cell membranes of HA-α_2_δ-1 expressed in tsA-201 cells. Top panel: α_2_δ-1 immunoblot, middle panel: HA immunoblot, both de-glycosylated to allow resolution between pro-α_2_δ-1 (upper band) and the cleaved form, α_2_-1 (lower band). Bottom panel: loading control endogenous GAPDH. The absolute % cleavage was 11.2 ± 1.0 % in control conditions. (E) Quantification of the effect of 1 and 100 µM SB-3CT on relative cleavage of α_2_δ-1, measured from HA immunoblots (normalized to that under control conditions). Data are mean ± SEM and individual data in 3 separate experiments, including that in (D), denoted by red, green and blue symbols). Statistical differences determined using 1-way ANOVA and Tukey post hoc test; *** *P* < 0.0001.

The α_2_δ subunits generally increase Ca^2+^ currents produced by Ca_V_α_1_/ß combinations, by a mechanism that is not yet completely understood (Dolphin, 2016; Kadurin et al., 2016). We have recently shown that α_2_δ-1 increases the density of Ca_V_2.2 channels inserted into the plasma membrane (Cassidy et al., 2014; Dahimene et al., 2018), and produces a large increase in calcium channel currents (Canti et al., 2005; Hendrich et al., 2008; Hoppa et al., 2012). For Ca_V_2.2, the interaction of Domain I extracellular loop 1 with the α_2_δ VWA domain is absolutely essential for its effect on trafficking and function (Canti et al., 2005; Dahimene et al., 2018).

In an extensive study, we have found that proteolytic processing of α_2_δ subunits is an essential step for activation of plasma membrane calcium channels. By replacing the proteolytic cleavage site in α_2_δ with an artificial site (α_2_(3C)δ), we found that uncleaved α_2_δ-1 inhibits native calcium currents in DRG neurons (Kadurin et al., 2016). Furthermore, uncleaved α_2_δ-1 inhibits presynaptic Ca^2+^ entry and vesicular release in hippocampal neurons (Ferron et al., 2018; Kadurin et al., 2016). We also showed that in non-neuronal cells the effect of α_2_δ on Ca_V_ channel activation can be separated from its trafficking role, in that uncleaved α_2_(3C)δ-1 and α_2_(3C)δ-3 can traffic Ca_V_2.2 channels to the plasma membrane, but these uncleaved constructs do not enhance Ca_V_2.2 currents, unless proteolytic cleavage is artificially induced (Kadurin et al., 2016). Thus, we proposed that proteolytic processing of α_2_δ subunits represents an activation step for calcium channel function, and pro-α_2_δ subunits maintain the channels in a state of low activation.

Upregulation of α_2_δ-1 protein is of importance in the development of neuropathic pain (Bauer et al., 2009; Luo et al., 2001; Newton et al., 2001; Patel et al., 2013), and α_2_δ-1 is also the drug target for gabapentinoid drugs used in neuropathic pain (Field et al., 2006). These drugs inhibit calcium channel trafficking when applied chronically (Cassidy et al., 2014; Hendrich et al., 2008). In the present study we have examined the nature of the enzyme(s) involved in proteolytic cleavage of α_2_δ subunits, since inhibition of its proteolytic cleavage could represent a novel point of therapeutic intervention.

## RESULTS

### Sequence of cleavage site in α_2_δ-1 predicts metalloproteases and ADAMs as candidate proteases

The proteolytic cleavage site in α_2_δ-1 has been identified to be between A and V in the sequence LEA∼VEME (De Jongh et al., 1990; Jay et al., 1991) (A945 and V946 in the rat sequence used here, Figure 1A), and we have shown that mutation of this site prevents the cleavage of α_2_δ-1 and abolishes the ability of α_2_δ-1 to increase calcium channel currents (Kadurin et al., 2016). Initial scrutiny of this sequence suggested a role for matrix metalloprotease (MMP) enzymes, specifically A Disintegrin and Metalloprotease (ADAM)10 or ADAM17/Tumor necrosis factor (TNF)-α converting enzyme (TACE). Although there are no absolute consensus motifs for proteolytic processing by these enzymes, there are preferred residues in the vicinity of the cleavage site (Caescu et al., 2009); for example the site in a well-established ADAM17 substrate Notch is IEA∼VKSE (Brou et al., 2000). However it should be noted that the proposed cleavage sites in α_2_δ-2 and α_2_δ-3 do not have similar primary sequences to that in α_2_δ-1 (Kadurin et al., 2016).

We have found previously that proteolytic cleavage of α_2_δ subunits is incomplete when it is expressed in cell lines, possibly attributable to saturation of the endogenous protease(s) required for cleavage (Davies et al., 2010; Kadurin et al., 2012; Kadurin et al., 2016). However, the degree of cleavage of α_2_δ-1 is increased at the plasma membrane and in detergent resistant membranes (DRMs), also called lipid rafts, to about 60% (Kadurin et al., 2012), and we found the same result in the present study. Importantly, the increased cleavage of α_2_δ-1 observed at the cell surface and in DRMs is not likely to be a result of differential trafficking of cleaved relative to uncleaved α_2_δ-1, since mutant uncleavable α_2_δ-1 is still able to reach the plasma membrane to the same extent as WT α_2_δ-1 (Kadurin et al., 2016).

### Chemical inhibitors of MMPs/ADAMs reduce proteolytic cleavage of α_2_δs

In order to examine whether MMPs or ADAMs were involved in α_2_δ-1 proteolytic cleavage, we first used a broad-spectrum hydroxamate metalloprotease inhibitor GM-6001, which inhibits both MMPs and ADAMs (Grobelny et al., 1992). GM-6001 produced more than 50% inhibition of α_2_δ-1 cleavage in whole cell membranes, when applied to tsA-201 cells at both 10 and 25 µM for 24 h (Figure 1B, C). The more selective inhibitor, SB-3CT, produced no inhibition at 1 µM, which is below the K_i_ for ADAM17 (∼4 µM) (Solomon et al., 2004), but resulted in about 40% reduction at 100 µM (*P* < 0.0001, Figure 1D, E). In contrast, the MMP 13 inhibitor CL-82198 (60 µM) produced no inhibition of α_2_δ-1 cleavage (Figure 1D).

### Reduced proteolytic cleavage of α_2_δ-1 in *ADAM17*^-/-^ but not *ADAM10*^-/-^ cell lines

The activation of endogenous MMPs and ADAMs often involves a complex sequential proteolytic cascade (Novak, 2004); for example, endogenous ADAM17 is activated by proteolytic cleavage with both furin and meprin ß (Wichert et al., 2019). For this and other reasons, overexpression of candidate proteases is often not a successful experimental route to identification of their role in biochemical pathways (see for example Romi et al., 2014). Thus, in order to examine the potential involvement of ADAM17 in cleavage of α_2_δ-1, we turned to HEK293 cell lines in which *ADAM10, ADAM17* or both protease genes were knocked out by CRISPR/Cas9 methodology, in comparison to the corresponding CRISPR wild type (WT) cells (Riethmueller et al., 2016).

In initial experiments we observed a marked reduction in proteolytic cleavage of α_2_δ-1 in whole cell lysates (WCL) of CRISPR *ADAM10*^-/-^/*ADAM17*^-/-^ double knockout cells, compared to CRISPR WT HEK293 cells (Figure 2A). This was also clearly observed in the cell surface-biotinylated fraction of these cells, in which greater basal cleavage is seen (Figure 2B). There was a 45.3 % reduction in proteolytic cleavage of α_2_δ-1 at the cell surface of *ADAM10*^-/-^/*ADAM17*^-/-^ cells (*P* = 0.0004; Figure 2C). Furthermore, similar results were obtained in DRMs from *ADAM10*^-/-^/*ADAM17*^-/-^ cells, (32 % reduction in α_2_δ-1 cleavage; *P* = 0.049; Figure S1A, B). There was no change in the distribution of α_2_δ-1 in DRMs from *ADAM10*^-/-^/*ADAM17*^-/-^ cells (Figure S1C).

**Figure 2:**
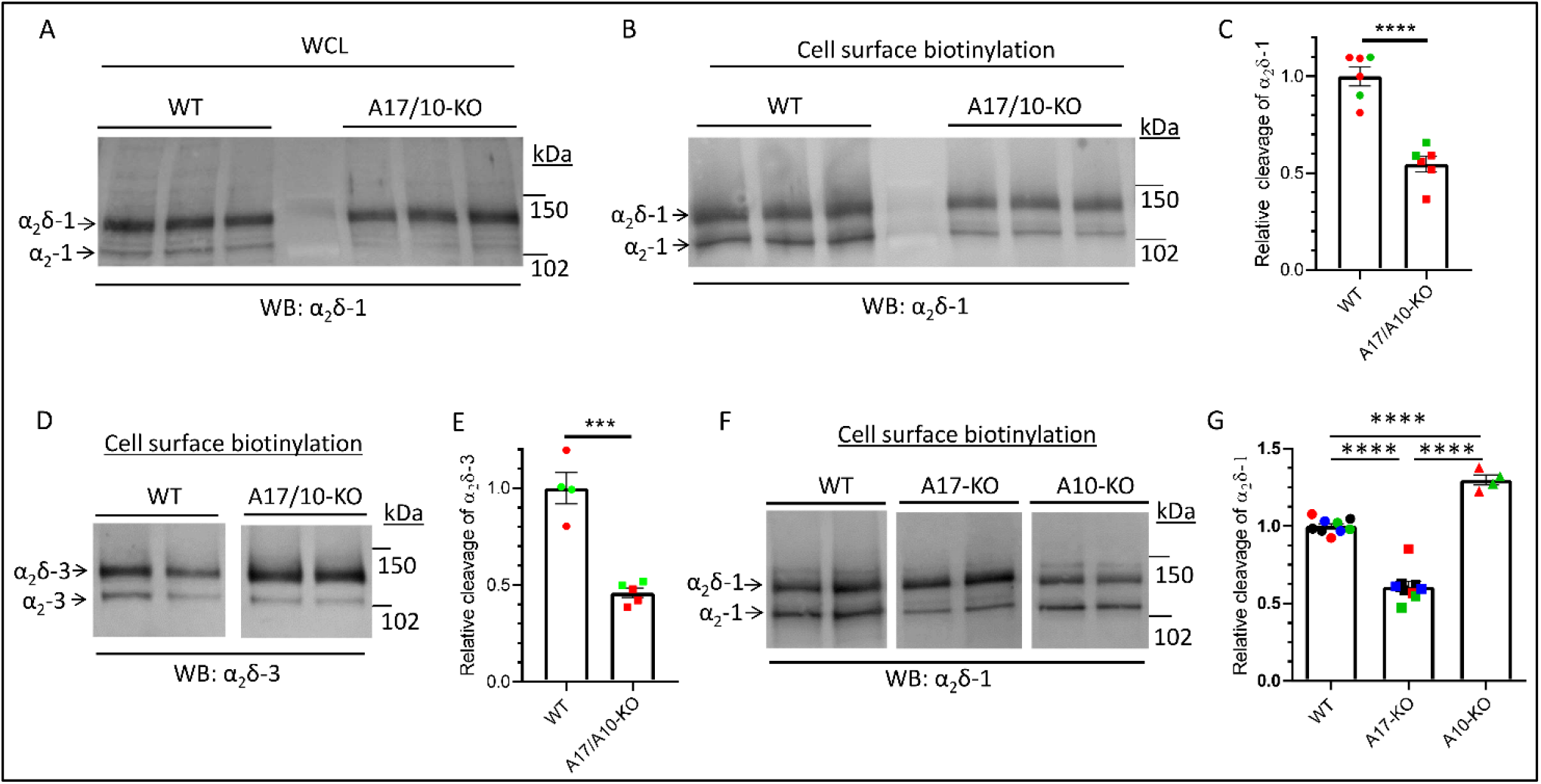
Cleavage of α_2_δ-1 and α_2_δ-3 is reduced in CRISPR *ADAM17*^-/-^ / *ADAM10*^-/-^ and *ADAM17*^-/-^ compared to CRISPR WT cells. (A) Effect of expression of α_2_δ-1 in CRISPR WT (lanes 1 - 3) compared to CRISPR *ADAM17*^-/-^ / *ADAM10*^-/-^ (A17/10-KO) HEK293 cells (lanes 4 - 6) on cleavage of HA-α_2_δ-1 (α_2_-1 immunoblot) in WCL (de-glycosylated to allow resolution between pro-α_2_δ-1 (upper band) and the cleaved form, α_2_-1 (lower band). The absolute % cleavage was 21.2 ± 1.4 % in CRISPR WT cells and 11.05 ± 1.2 % in *ADAM17*^-/-^ / *ADAM10*^-/-^ cells (n = 4 experiments). (B) Effect of expression of α_2_δ-1 in CRISPR WT (lanes 1 - 3) compared to *ADAM17*^-/-^ / *ADAM10*^-/-^ (A17/10-KO) HEK293 cells (lanes 4 - 6) on cleavage of HA-α_2_δ-1 (α_2_-1 immunoblot) in deglycosylated cell surface biotinylated fractions. The absolute % cleavage was 37.4 ± 2.4 % in CRISPR WT cells (n = 6 experiments). (C) Quantification of the effect of expression in *ADAM17*^-/-^ / *ADAM10*^-/-^ cells on relative cleavage of α_2_δ-1 in cell surface biotinylated fraction, normalized to that in CRISPR WT cells. Data are mean ± SEM and individual data from 6 separate experiments, performed on two different transfections (red and green symbols). Statistical difference determined using Student’s t test; *P* = 0.0004. (D) Effect of expression of α_2_δ-3 in CRISPR WT (lanes 1, 2) compared to *ADAM17*^-/-^ / *ADAM10*^-/-^ HEK293 cells (lanes 3, 4) on cleavage of HA-α_2_δ-3 in cell surface biotinylated fraction (α_2_-3 immunoblot), de-glycosylated to allow resolution between pro-α_2_δ-3 (upper band) and the cleaved form, α_2_-3 (lower band). The absolute % cleavage was 23.3 ± 2.5 % in CRISPR WT cells (n = 4). (E) Quantification of the effect of *ADAM17*^-/-^ / *ADAM10*^-/-^ on relative cleavage of α_2_δ-3 in cell surface biotinylated fraction (normalized to that in CRISPR WT cells). Data are mean ± SEM and individual data in 4-5 separate experiments, performed on two separate transfections (red, and green symbols). Statistical difference determined using Student’s t test; *** *P* = 0.0002). (F) Effect of expression of α_2_δ-1 in CRISPR WT (lanes 1, 2) compared to *ADAM17*^-/-^ (lanes 3, 4) and *ADAM10*^-/-^ (lanes 5, 6) HEK293 cells on cleavage of HA-α_2_δ-1 (α_2_-1 immunoblot) in cell surface biotinylated fractions, de-glycosylated to allow resolution between pro-α_2_δ-1 (upper band) and the cleaved form, α_2_-1 (lower band). The absolute % cleavage was 44.5 ± 2.7 % in CRISPR WT cells (n = 9). (G) Quantification of the effect of expression in *ADAM10*^-/-^ and *ADAM17*^-/-^ cells on relative cleavage of α_2_δ-1 in cell surface biotinylated fraction, normalized to that in CRISPR WT cells. Data are mean ± SEM and individual data for 9 separate experiments from four different transfections (all including WT and *ADAM17*^-/-^ cells, and 4 also including *ADAM10*^-/-^cells; coloured symbols refer to different experiments). Statistical differences determined using one-way ANOVA and Tukey’s multiple comparison test; **** *P* < 0.0001.

Although the identified α_2_δ-3 proteolytic cleavage motif has a primary sequence that is different from that of α_2_δ-1 (Kadurin et al., 2016), the ADAM proteases support a wide divergence of cleavage motifs (see for example Chen et al., 2020). We therefore performed the same experiment, using α_2_δ-3 as substrate, and observed a similar reduction in its proteolytic cleavage of 53.9 %, in the cell-surface biotinylated fraction of *ADAM10*^-/-^/*ADAM17*^-/-^ cells compared to CRISPR WT cells (*P* = 0.0002, Figure 2D, E).

Next, we examined whether the loss of ADAM10 or ADAM17 was responsible for this effect, by using single knockout cell lines. We found a significant 39.1 ± 3.4 % (*P* < 0.0001) reduction in proteolytic cleavage of α_2_δ-1 in cell surface-biotinylated fractions from *ADAM17*^-/-^ compared to control cells, whereas there was a small increase in α_2_δ-1 cleavage in *ADAM10*^-/-^ cells (Figure 2F, G).

### Effect of *ADAM17* or *ADAM10* knockout on calcium channel currents and cell surface expression

We then wished to examine whether the reduction in proteolytic cleavage of α_2_δ-1 by ADAM17 had a functional effect on Ca_V_ currents, as would be predicted from our previous study, in which we showed non-cleavable α_2_δ constructs were non-functional in this regard (Kadurin et al., 2016). We first examined Ca_V_ currents formed by Ca_V_2.2 together with β1b and α_2_δ-1, expressed in *ADAM17*^-/-^ or *ADAM10*^-/-^ HEK293 cells, compared to CRISPR WT cells. There was a clear reduction in I_Ba_ (by 44.5 % at +5 mV, *P* = 0.0002, Figure 3A, B) in *ADAM17*^-/-^ but not *ADAM10*^-/-^ cells, with no change in the potential for half-activation, V_50, act_ (Figure 3B).

**Figure 3:**
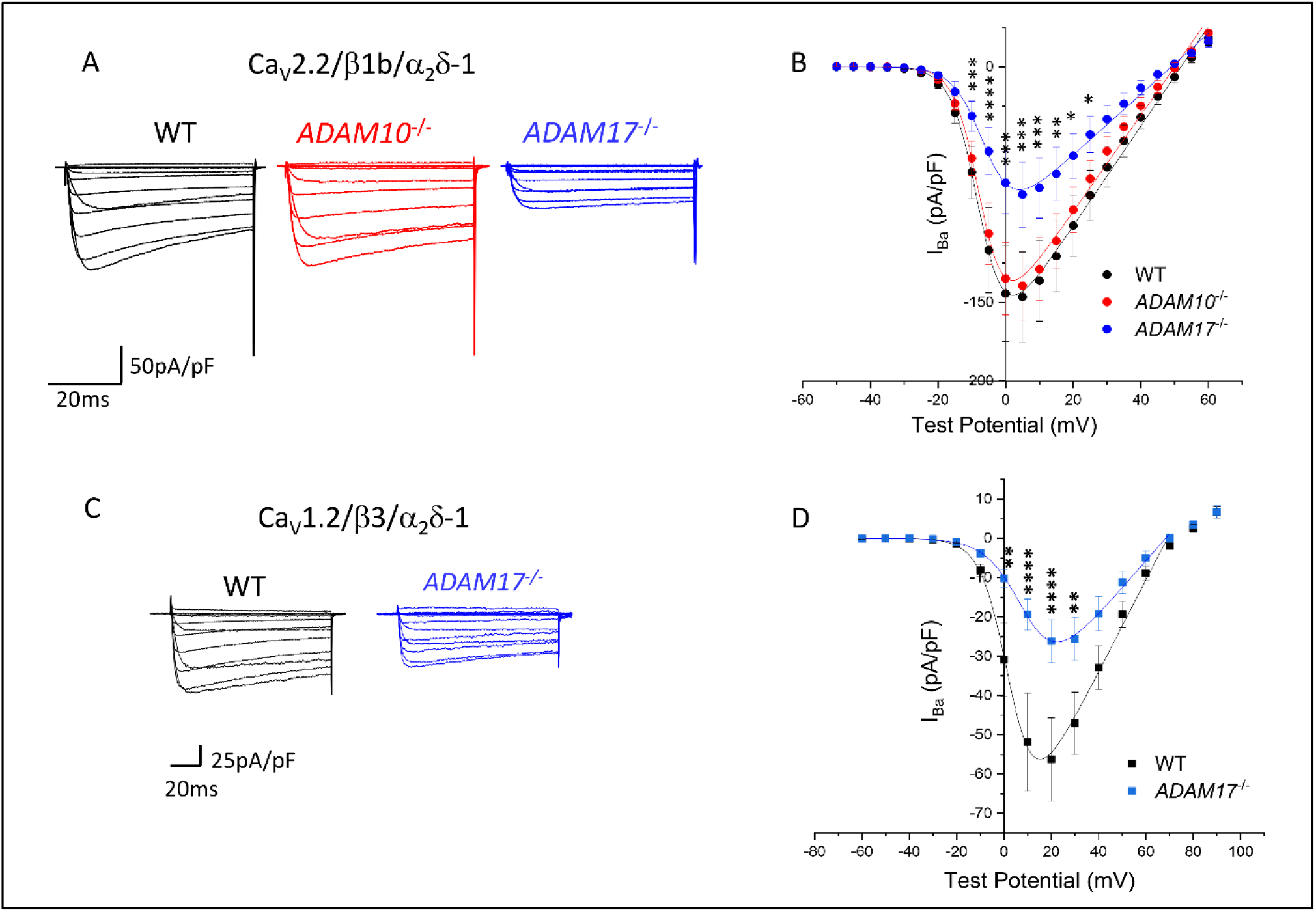
Knockout of *ADAM17*, but not *ADAM10*, decreases Ca_V_2.2 and Ca_V_1.2 currents. (A) Examples of I_Ba_ currents for Ca_V_2.2 expressed in CRISPR WT HEK293 cells (black), *ADAM10*^-/-^ cells (red) and *ADAM17*^-/-^ cells (blue). Ca_V_2.2 is co-expressed with β1b and α_2_δ-1. Holding potential −80 mV, steps between −50 mV and +60 mV for 50 ms. (B) Mean (± SEM) current-voltage (*I-V*) relationships for the conditions shown in (A). Control (black; n = 23), *ADAM10*^-/-^ (red, n =29) and *ADAM17*^-/-^ (blue; n = 21). The individual and mean *I-V* data were fit with a modified Boltzmann equation (see Methods). *V*_50, act_ values are −5.9 ± 1.4 mV (Control), −5.8 ± 0.8 mV (*ADAM10*^-/-^) and −2.5 ± 1.36 mV (*ADAM17*^-/-^). Two-Way ANOVA with Sidak’s *post hoc* test correction for multiple comparisons was performed for the *I-V P* < 0.05, *P* < 0.01, *P* < 0.001, *P* < 0.0001. (C) Examples of I_Ba_ currents for Ca_V_1.2 expressed in CRISPR WT HEK293 cells (black) and *ADAM17*^-/-^ cells (blue). Ca_V_1.2 is co-expressed with β3 and α_2_δ-1. Holding potential −80 mV, steps between −60 and +90 mV for 50 ms. (D) Mean (± SEM) *I-V* relationships for the conditions shown in (C). Control (black; n = 26) and *ADAM17*^-/-^ (blue; n = 19). The individual and mean *I-V* data were fit with a modified Boltzmann equation as in (B). *V*_50, act_ values are 5.8 ± 1.0 mV (Control) and 6.5 ± 1.9 mV (*ADAM17*^-/-^). Statistical differences between the two sets of *I-V* were examined at each potential and corrected for multiple t tests with Holm Sidak’s *post hoc* correction *P* < 0.01, *P* < 0.0001.

If the reduction in Ca_V_2.2-mediated I_Ba_ in *ADAM17*^-/-^ cells relates to the reduced cleavage of α_2_δ-1, then it should also occur for another channel subtype. We therefore examined Ca_V_1.2, co-expressing it with α_2_δ-1 and a different β (β3), comparing CRISPR WT cells with *ADAM17*^-/-^ cells (Figure 3C, D). We found a reduction in peak I_Ba_ for Ca_V_1.2 in ADAM17^-/-^ cells (by 53.5 % at +20 mV, P < 0.0001), which was similar to that found for Ca_V_2.2, thereby implicating α_2_δ-1 in this reduction. As for Ca_V_2.2, there was no significant change in the V_50, act_ (Figure 3D).

In order to determine whether the effect on Ca_V_ currents of expression in *ADAM17*^-/-^ cells was related to an effect on trafficking of the channels, we examined the cell surface expression of Ca_V_2.2 (in the presence of β1b and α_2_δ-1), using the exofacial HA epitope on a GFP_Ca_V_2.2-HA construct, relative to its cytoplasmic expression, measured by GFP, as described previously (Dahimene et al., 2018; Meyer et al., 2019). In contrast to the marked reduction in Ca_V_2.2 currents in *ADAM17*^*-/*-^ cells, there was no effect on cell surface expression of the channel, as measured by the HA/GFP ratio (Figure 4A, B), indicating that there was no influence of *ADAM17* knockout on Ca_V_2.2 trafficking. This would agree with our previous finding that Ca_V_2.2 cell surface expression in non-neuronal cells was still increased by a non-cleavable α_2_δ-1 construct (Kadurin et al., 2016). This result reinforces our finding that the cleavage of α_2_δ-1 is not essential for calcium channel trafficking to the plasma membrane in undifferentiated cell lines, but is essential for enhancing calcium channel function (Kadurin et al., 2016). Of interest, there was a small increase in Ca_V_2.2 cell surface expression in *ADAM10*^-/-^ cells, relative to the CRISPR WT cells (Figure 4B), and this could relate to the increased proteolytic cleavage of α_2_δ-1 in the cell surface fraction of *ADAM10*^-/-^ cells (Figure 2G), since the % cleavage of α_2_δ-1 is elevated in cell surface biotinylated fractions (Kadurin et al., 2012).

**Figure 4:**
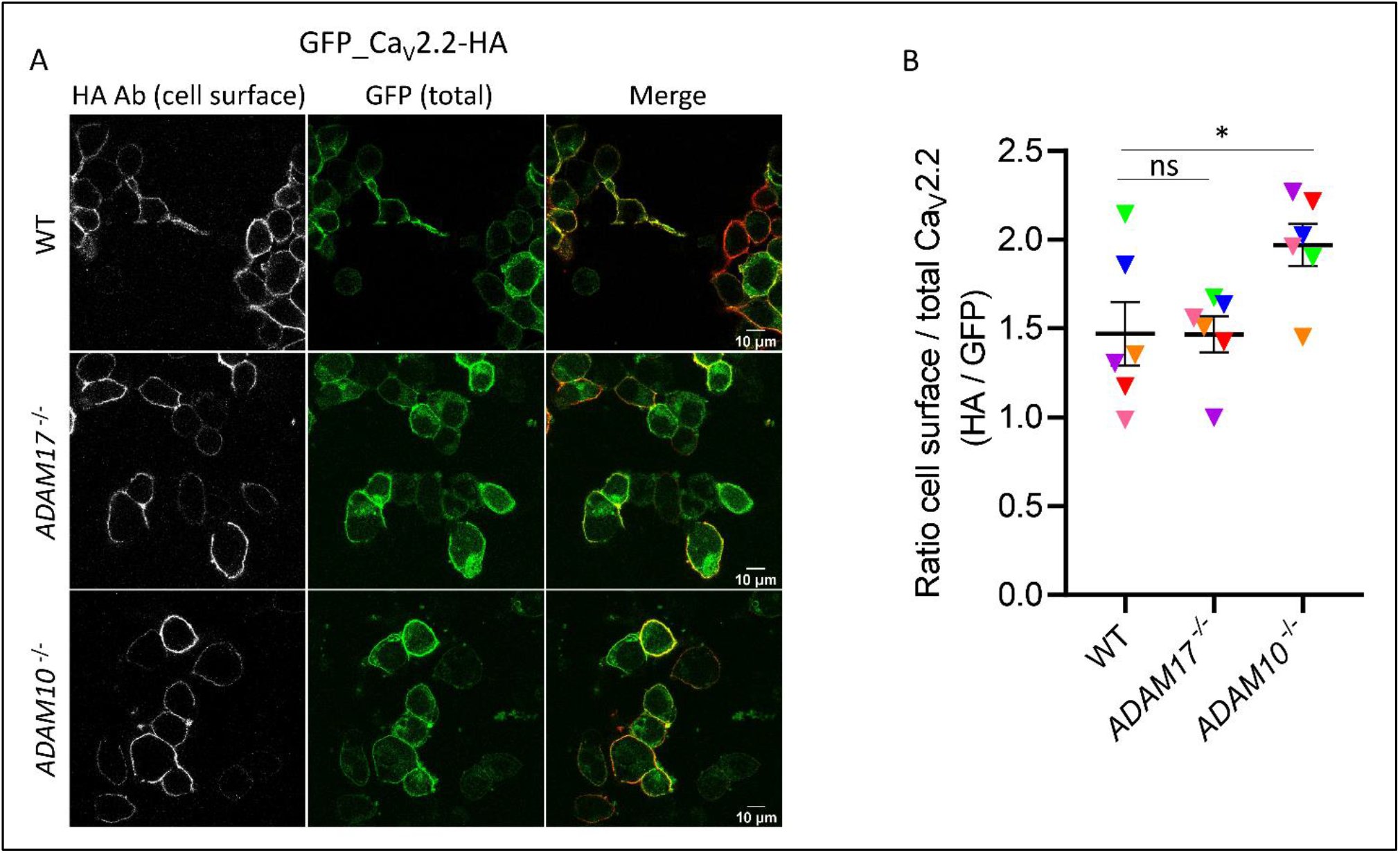
*ADAM17* knockout does not alter the cell surface expression of Ca_V_2.2 at the plasma membrane. (A) Confocal images of GFP_Ca_V_2.2-HA expressed in CRISPR WT (top), *ADAM17*^-/-^ (middle) or *ADAM10*^-/-^ (bottom) HEK293 cells. All conditions contained β1b and α_2_δ-1. Cell surface staining of GFP_Ca_V_2.2-HA was obtained by incubating the cells with HA Ab in non-permeabilized cells (left, white). The total expression of GFP_Ca_V_2.2-HA is determined by the cytoplasmic GFP signal (middle), the merged images are shown on the right (HA in red). Scale bars, 10 μm. (B) Scatter plot (mean ± SEM with individual data points for six independent transfections, with all conditions in parallel), showing the ratio of cell surface to total Ca_V_2.2 (HA / GFP) in CRISPR WT, *ADAM17*^-/-^or *ADAM10*^-/-^ HEK293 cells. Each individual data point represents a mean of ratio HA / GFP of ∼100 cells/experiment. ns = non-significant for WT vs *ADAM17*^-/-^; *, *P* = 0.0379 for WT vs *ADAM10*^-/-^, one-way ANOVA and Tukey’s *post hoc* test, correcting for multiple comparisons).

### Subcellular site of proteolytic cleavage

In a previous study we found that cleaved α_2_δ-1 is associated with a mature glycosylation pattern, as N-linked glycans are trimmed and modified in the Golgi apparatus (Aebi et al., 2010), although some membrane proteins can bypass this route (Hanus et al., 2016). Conversely, uncleaved α_2_δ-1 primarily possesses immature endoplasmic reticulum (ER)-associated glycosylation that can be removed by endoglycosidase-H (Endo-H) in WCL fractions (Kadurin et al., 2017) (see diagram in Figure 5A). This suggests that proteolytic cleavage is likely to be associated mainly with post-ER organelles, including the Golgi apparatus (Kadurin et al., 2017). Nevertheless, it is also the case that α_2_δ cleavage can be induced to occur on the plasma membrane (Kadurin et al., 2016).

**Figure 5:**
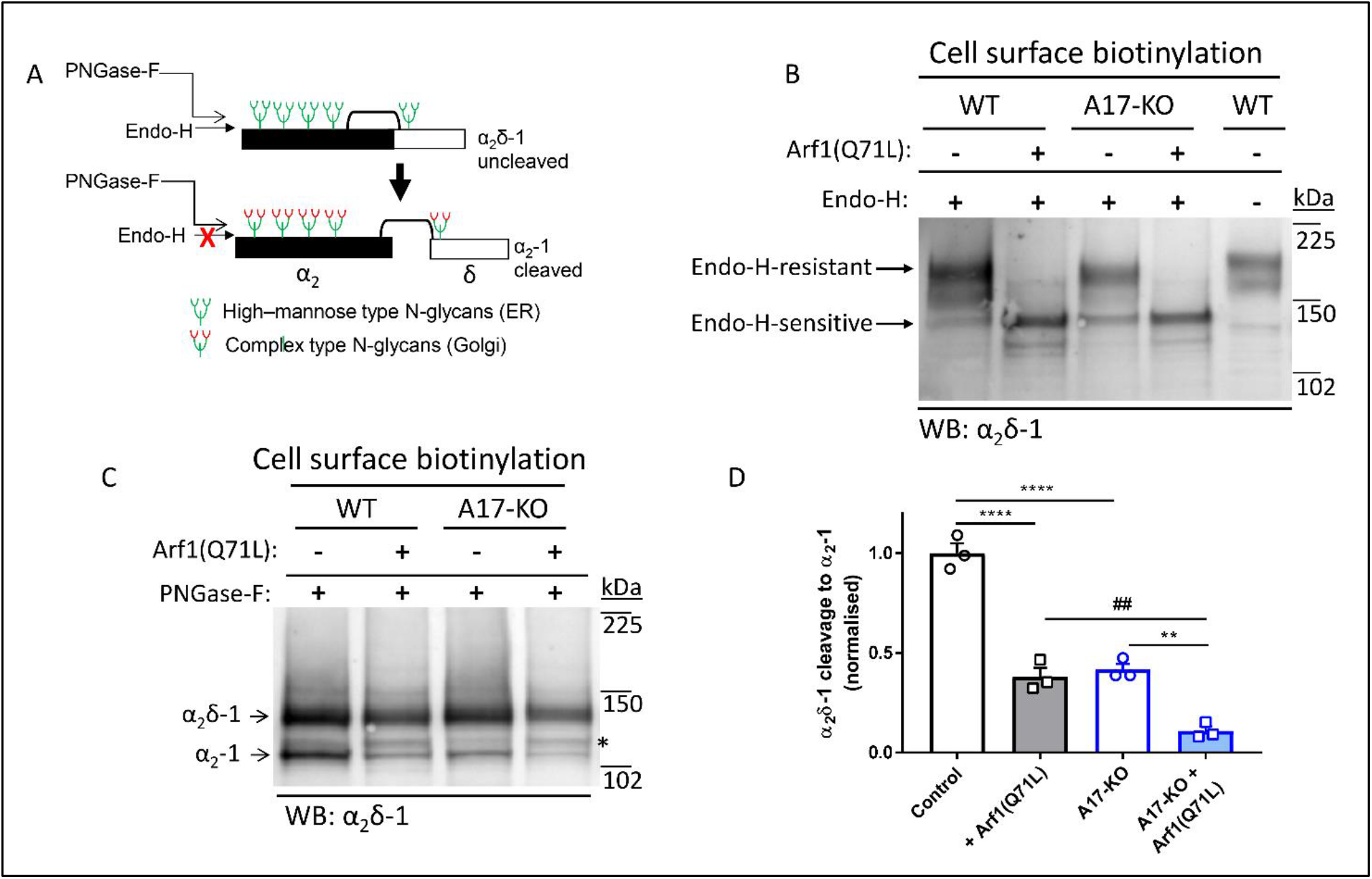
Effect of block of ER-Golgi trafficking and stimulation of alternative route to cell surface by Arf1(Q71L) on cleavage and N-glycosylation pattern of α_2_δ-1 in CRISPR WT and *ADAM17*^-/-^ cells. (A) Schematic representation of post-translational modifications of α_2_δ-1. Immature glycosylation, occurring in the ER, is sensitive to Endo-H, whereas mature glycosylation, occurring in the Golgi, is resistant to Endo-H. There are differential effects of Endo-H on uncleaved (top) and cleaved (bottom) α_2_δ-1. (B) Effect of expression of α_2_δ-1 in CRISPR WT (lanes 1, 2, 5) and *ADAM17*^-/-^ (A17-KO) cells (lanes 3, 4) in the absence (lanes 1, 3, 5) and presence (lanes 2, 4) of the ER-to-Golgi blocker, Arf1(Q71L), in cell surface biotinylated samples, to show fraction at the plasma membrane, either treated with Endo-H (lanes 1 - 4) or left untreated (lane 5) for comparison. The sizes of the Endo-H-resistant bands (after mature glycosylation in the Golgi) and Endo-H-sensitive bands (after blocking ER-to-Golgi transport using Arf1(Q71L)) are indicated with arrows. Representative of n = 3 separate experiments. (C) Effect of expression of α_2_δ-1 in CRISPR WT (lanes 1, 2) and *ADAM17*^-/-^ (A17-KO) cells (lanes 3, 4) in the absence (lanes 1, 3) and presence (lanes 2, 4) of Arf1(Q71L). Samples are biotinylated and fully deglycosylated with PNGase-F to show uncleaved-α_2_δ-1 (upper band) and cleaved α_2_-1 (lower band). The absolute % cleavage of α_2_δ-1 to cleaved α_2_-1 was 27.3 ± 1.3 % in control CRISPR WT cells (n = 3). * indicates an intermediate species, which may represent cleavage of α_2_δ-1 at an alternative site, or an intermediate product. (D) Quantification of the effect of of Arf1(Q71L) (shaded compared to open bars) in WT (black bars) and *ADAM17*^-/-^ cells (blue bars) on cleavage of cell surface biotinylated α_2_δ-1, normalized to that in control CRISPR WT cells. Data are mean ± SEM and individual data for 3 separate experiments, including that in (C). Statistical differences determined using one-way ANOVA and Sidak’s multiple comparison test; **** *P* < 0.0001; ** *P* = 0.0017; ## *P* = 0.0037.

In order to examine whether α_2_δ-1 needs to be trafficked through the Golgi to be proteolytically cleaved by ADAM17 protease, we pursued several experimental routes. Firstly, we used a constitutively-active mutant ADP ribosylation factor (Arf)1 (Q71L), which blocks traffic between ER and Golgi (Dascher and Balch, 1994), and promotes the utilization of an unconventional endosomal pathway to the cell surface that bypasses the Golgi apparatus (Gee et al., 2011). Confirming this alternative trafficking route, we found that, in the presence of Arf1(Q71L), α_2_δ-1 was still able to reach the cell surface (Figure 5B).

Under control conditions, α_2_δ-1 in the cell surface-biotinylated fraction was mainly Endo-H-resistant in both CRISPR WT and *ADAM17*^-/-^ cells, indicating that it had been trafficked to the plasma membrane via the Golgi apparatus, where it had obtained mature N-glycans (Figure 5B, lanes 1, 3). By contrast, in the presence of Arf1(Q71L), α_2_δ-1 in the cell surface-biotinylated fraction was completely Endo-H-sensitive in both CRISPR WT and *ADAM17*^-/-^ HEK293 cells (Figure 5B, lanes 2, 4), indicating that, in this case, α_2_δ-1 at the plasma membrane contained only immature N-glycans derived from the ER, and that it had not been processed in the Golgi. As expected, in the WCL most α_2_δ-1 was Endo-H-sensitive, suggesting it was derived from the ER (Figure S2).

In agreement with data presented in Figure 2, we observed less proteolytic cleavage of cell surface-biotinylated α_2_δ-1 to α_2_-1 in *ADAM17*^*-/-*^ compared to WT cells (58.3 % reduction, Figure 5C, D). Proteolytic cleavage of cell surface α_2_δ-1 was also significantly reduced by Arf1(Q71L) expression in WT cells by 61.9 %, and residual cleavage was further reduced in *ADAM17*^*-/-*^ cells by 73.9 % (Figure 5C, D), indicating that the Golgi apparatus is an important site of proteolytic cleavage of α_2_δ-1. Interestingly expression of Arf1(Q71L) also promoted the appearance of an intermediate MW species of cleaved α_2_δ-1, which may represent cleavage of α_2_δ-1 at an alternative site, or an intermediate product (Figure 5C, *).

The conclusion that cleavage of α_2_δ proteins is associated in part with the Golgi apparatus was also borne out by subcellular fractionation of an α_2_δ-3 stable SH-SY5Y cell line, in which uncleaved α_2_δ-3 is associated with an ER marker, protein disulfide isomerase (PDI), whereas the appearance of cleaved α_2_-3 and δ-3 moieties are associated with the presence of the Golgi marker, g97 (Figure S3A, B).

In order to examine whether cleavage of α_2_δ proteins could also occur on the cell surface, we examined the efficacy of the tissue inhibitor of metalloproteases (TIMP-3). TIMPs are endogenously expressed small proteins, secreted in the extracellular matrix, which differentially inhibit particular MMPs and ADAMs (Brew and Nagase, 2010), although their actions are complex and may also involve activation of some MMPs via ternary complex formation (Zhao et al., 2004). TIMP-3 inhibits most MMPs and ADAMs, including ADAM17 (Brew and Nagase, 2010), whereas the catalytically active N-terminal domain (N-TIMP-3) selectively inhibits ADAM17 (Brew and Nagase, 2010), but not ADAM10 (Rapti et al., 2008). Thus, for this study we applied N-TIMP-3 protein (100 nM, for 24 h) extracellularly to cells expressing α_2_δ-1. As expected, N-TIMP-3 did not inhibit cleavage of α_2_δ-1 in the WCL (Figure 6A, B), but it significantly reduced cleavage of cell surface α_2_δ-1 (by 21%, *P* = 0.023; Figure 6A, C). This result indicates that at least some cleavage of α_2_δ-1 can occur at the plasma membrane. This experiment also confirms that ADAM17 is likely to be involved in this process.

**Figure 6:**
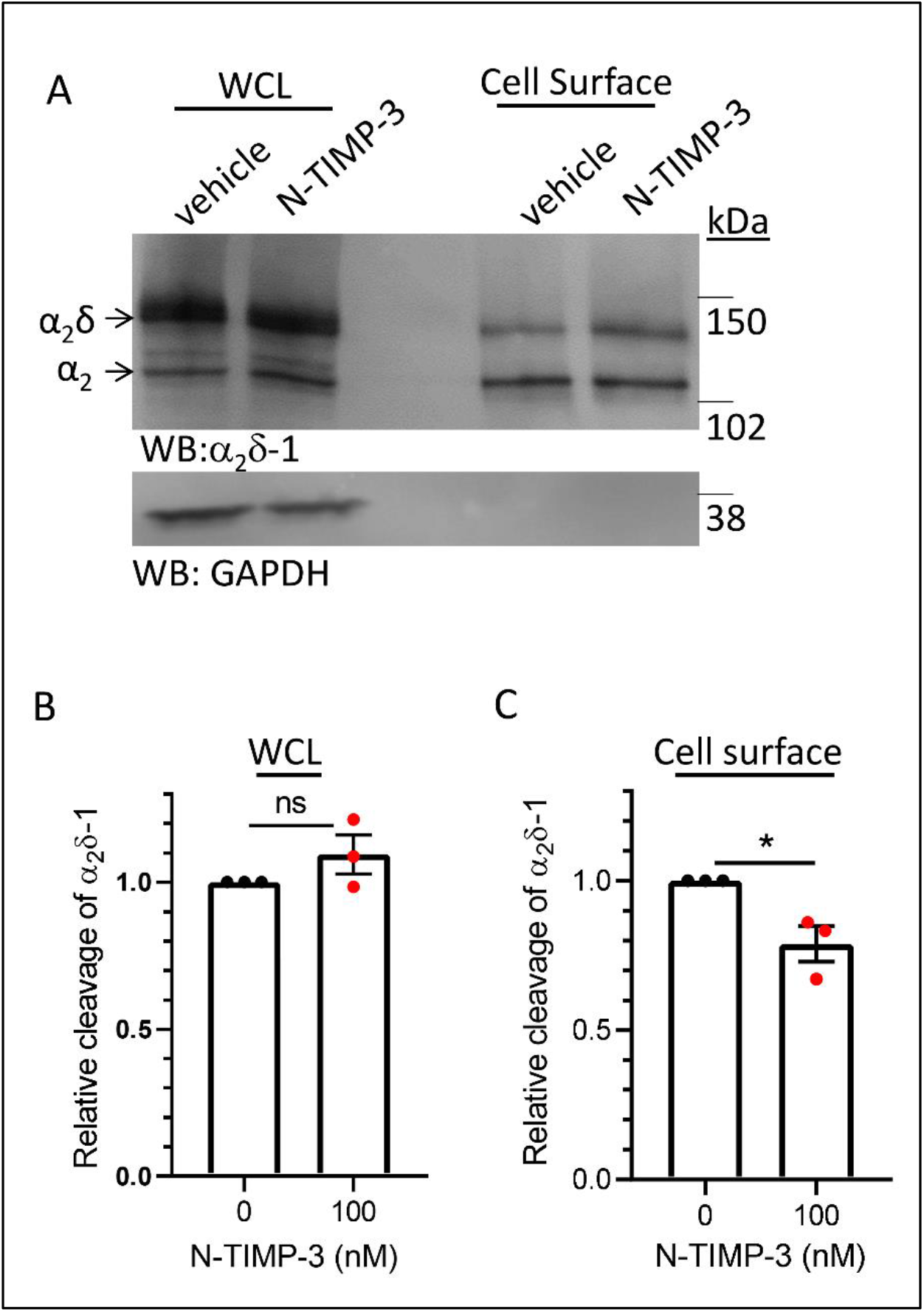
N-TIMP-3 inhibits cleavage of α_2_δ-1 on cell surface of tsA-201 cells. (A) Effect of extracellular application of N-TIMP-3 (lanes 2, 5) compared to control (lanes 1, 4), on cleavage in WCL (lanes 1, 2) and cell surface biotinylated fractions (lanes 4, 5) of HA-α_2_δ-1 expressed in tsA-201 cells (upper panel: α_2_δ-1 immunoblot), de-glycosylated to allow resolution between pro-α_2_δ-1 (upper band) and the cleaved form, α_2_-1 (lower band). Lower panel, loading and biotinylation control: endogenous GAPDH. (B, C) Quantification of the effect of N-TIMP-3 application on relative cleavage of α_2_δ-1 in WCL (B) and at plasma membrane (C) (mean ± SEM and individual data shown for 3 separate experiments, each normalized to that under control conditions). * *P* = 0.0231 (Student’s t test).

## DISCUSSION

### Cleavage of α_2_δ-1 involves ADAM17

The importance of α_2_δ-1 in multiple disorders (Dolphin, 2014), and its relevance as a therapeutic target (Field et al., 2006), coupled with our finding that its proteolytic cleavage into mature disulfide-bonded α_2_ and δ is required for the enhancement of calcium channel function (Kadurin et al., 2016), suggested to us that targeting the proteolytic cleavage of α_2_δ-1 could represent a novel site for therapeutic intervention. The structure of the α_2_δ-1 subunit (Wu et al., 2016) does not completely resolve the proteolytic cleavage site, and the C-terminus of α_2_ and N-terminus of δ are not co-located, with some missing sequence. This might indicate that either these termini are dynamic within the structure, and a conformational change might occur on proteolytic cleavage of α_2_δ such that they could move apart, or that a loop of α_2_δ-1 might be excised by successive proteolytic cleavages, as occurs for some other proteins (Wichert et al., 2019).

In this study we have identified that ADAM17 is a key protease involved in proteolytic processing of pro-α_2_δ-1 and α_2_δ-3 subunits. We provide three lines of evidence for the role of ADAM17: firstly, proteolytic cleavage is inhibited by chemical inhibitors of particular MMPs and ADAMs, including ADAM17. Secondly, proteolytic cleavage of α_2_δ-1 and α_2_δ-3 is markedly reduced in cell lines by knockout of *ADAM17* but not by knockout of *ADAM10*. Thirdly, proteolytic cleavage of α_2_δ-1 is reduced by the N-terminal active domain of TIMP-3 (N-TIMP-3), which selectively inhibits ADAM17. Nevertheless, the incomplete block of cleavage of α_2_δ-1 in ADAM17^-/-^ cells indicates other proteases are likely to be involved, either at the same or nearby cleavage sites between α_2_ and δ, as also occurs for many other proteins, for example amyloid precursor protein (Eggert et al., 2018). This is borne out by the observation of an additional higher molecular weight α_2_ band observed here. Furthermore, ADAM17 is itself involved in protease cascades, and requires activation by other proteases (Wichert et al., 2019; Zunke and Rose-John, 2017).

### The importance of α_2_δ-1 cleavage for calcium channel function

In heterologous systems, the co-expression of α_2_δ subunits has been shown by many groups to increase the recorded calcium channel currents for Ca_V_1 and Ca_V_2 channels by 3 - 10-fold (Barclay et al., 2001; Canti et al., 2005; Gao et al., 2000; Hendrich et al., 2008; Hobom et al., 2000; Klugbauer et al., 1999; Mori et al., 1991). We found previously that this enhancement requires the proteolytic cleavage of α_2_δ into α_2_ and δ, and for Ca_V_2.2 we further showed, using uncleavable α_2_δ constructs, that this process was independent of the increased cell surface expression of the channel (Kadurin et al., 2016). Thus, the effect of α_2_δ-1 and α_2_δ-3 on calcium channel trafficking to the plasma membrane in undifferentiated cell lines did not require α_2_δ proteolytic cleavage; nevertheless the α_2_δ-mediated potentiation of calcium channel currents requires a molecular switch provided by proteolytic cleavage of the α_2_δ subunit (Kadurin et al., 2016).

Here we show that knockout of *ADAM17*, which inhibits α_2_δ-1 cleavage, has a clear functional consequence for calcium channel currents, since both Ca_V_2.2 and Ca_V_1.2 currents (both in the presence of α_2_δ-1, but with different β subunits, β1b and β3, respectively) are significantly reduced by 40 – 50 % in *ADAM17*^-/-^ cells, compared to either CRISPR WT or *ADAM10*^-/-^ cells. In contrast, and as also predicted from our previous study (Kadurin et al., 2016), there was no effect of *ADAM17* knockout on the α_2_δ-1-mediated increase in cell surface expression of Ca_V_2.2.

### Subcellular site of proteolytic processing of α_2_δ

In a previous study we found that native α_2_δ-1 was completely processed in axons (Kadurin et al., 2016), despite being intracellular until it reached synaptic terminals (Bauer et al., 2009). This agrees with our present results indicating that α_2_δ processing begins to occur intracellularly in the Golgi complex, and also with the finding that uncleaved α_2_δ, unlike cleaved α_2_δ, exhibits an immature glycosylation pattern (Kadurin et al., 2017).

This conclusion is supported by our finding that mutant Arf1(Q71L), which promotes an alternative pathway for α_2_δ-1 to reach the cell surface, bypassing the Golgi apparatus (Hanus et al., 2016), reduces proteolytic cleavage of α_2_δ-1, in synergy with *ADAM17* knockout. However, a proportion of α_2_δ-1 is still cleaved in the presence of Arf1(Q71L), despite it being completely Endo-H-sensitive, suggesting that the Golgi is not the only site where cleavage can occur, and that mature N-glycosylation is not essential for proteolytic cleavage to occur. Furthermore, the finding that extracellular application of N-TIMP-3 protein produces some inhibition of α_2_δ-1 cleavage indicates that cleavage can also occur on the cell surface, although it should be noted that TIMP proteins can also be endocytosed (Fan and Kassiri, 2020), and thus cleavage of α_2_δ-1 could also occur in the endosomal network. The subcellular distribution of ADAM17 would agree with these findings as although most ADAM17 is present intracellularly (Lorenzen et al., 2016), and it is activated by furin in the Golgi complex (Endres et al., 2003), nevertheless some active ADAM17 is expressed on the cell surface (Lorenzen et al., 2016).

### Relevance of α_2_δ-1 function to pain

There is strong upregulation of α_2_δ-1 mRNA and protein in rodent neuropathic injury models (Bauer et al., 2009; Luo et al., 2001; Newton et al., 2001). Furthermore, overexpression of α_2_δ-1 mimics neuropathic allodynia (Li et al., 2006), whereas knockout of α_2_δ-1 markedly delays the onset of neuropathic mechanical allodynia (Patel et al., 2013). We have shown previously that native pro-α_2_δ-1 (presumably newly synthesized) can be observed in the cell bodies of dorsal root ganglion neurons (Kadurin et al., 2016). ADAM17 has many substrates (Zunke and Rose-John, 2017) and ADAM17 inhibitors have many potential therapeutic targets (Arribas and Esselens, 2009). However, *ADAM17* knockout mice are non-viable (Peschon et al., 1998), which has hampered research into its many functions. Nevertheless, partial knockdown of ADAM17 has been shown to impair mechanical, heat, and cold nociception (Quarta et al., 2019), although the mechanism for this was not explored. Furthermore, *ADAM17* knockout in specific neurons has been found to reduce excitation (Xu et al., 2019).

## Conclusion

We know from our previous work that proteolytic cleavage into mature α_2_δ is essential for the enhancement of Ca_V_ function (Ferron et al., 2018; Kadurin et al., 2016). Our present study identifies a key protease involved in proteolytic maturation of α_2_δ-1 and α_2_δ-3 to be ADAM17, and in agreement with this, knockout of *ADAM17* inhibited the ability of α_2_δ-1 to enhance calcium currents. Coupled with our finding that some proteolytic cleavage of α_2_δ-1 can occur at the plasma membrane, this opens a potential novel therapeutic target, for example in neuropathic pain.

## CONTACT FOR REAGENTS

Further information and requests for resources and reagents should be directed to and will be fulfilled where possible by the Lead Contacts, Annette Dolphin and Ivan Kadurin, Department of Neuroscience, Physiology and Pharmacology, University College London, Gower Street, London, WC1E 6BT, UK Tel: +44-20-7679 3276

## METHODS

### Molecular biology

The following cDNAs were used: Ca_V_2.2 (rabbit, D14157), Ca_V_2.2-HA (Cassidy et al., 2014), GFP_Ca_V_2.2-HA (Macabuag and Dolphin, 2015), Ca_V_1.2 (rat; M67515.1), β1b (rat, X61394) (Pragnell et al., 1991), β3 (rat; M88751), α_2_δ-1 (rat, M86621) (Kim et al., 1992), HA-α_2_δ-1 (Kadurin et al., 2012), α_2_δ-3 (AJ010949), HA-α_2_δ-3 (Kadurin et al., 2016), mCherry (Shaner et al., 2004), mut2-GFP (Cormack et al., 1996), Arf(Q71L)-CFP (Addgene plasmid # 128149) (Presley et al., 2002), CFP replaced with mCherry. The cDNAs were in the pcDNA3 vector for expression in tsA-201 and HEK293 cells. CD8 cDNA (Rougier et al., 2005) was included as a transfection marker where stated.

### Antibodies and other materials

Antibodies (Abs) used were: anti-α_2_δ-1 Ab (mouse monoclonal, Sigma-Aldrich), anti-α_2_δ-3 and anti-δ-3 Ab (Davies et al., 2010), Ab anti-HA Ab (rat monoclonal, Roche), anti-GAPDH Ab (mouse monoclonal, Ambion), anti-FLAG Ab (rabbit polyclonal; Sigma), anti-PDI (mouse monoclonal, Ambion), anti-g97 (rabbit polyclonal; Abcam), anti-flotillin Ab monoclonal, BD Biosciences). For immunoblotting, secondary Abs (1:2000) were anti-rabbit-Horseradish Peroxidase (HRP), and anti-mouse HRP (Biorad). For immunocytochemistry, anti-rat-Alexa Fluor 594 was used at 1/500 (ThermoFisher).

The metalloprotease inhibitors GM6001 (BML-EI300, Enzo Life Sciences), SB-3CT (BMEI325, Enzo Life Sciences) and MMP-13 inhibitor (BML-EI302, Enzo Life Sciences) were dissolved in DMSO (or water for MMP-13 inhibitor) and used at the concentrations stated. N-TIMP-3 protein (expressed in *E. coli* as previously described (Kashiwagi et al., 2001), or control samples in the absence of N-TIMP-3, were pre-incubated with heparin (200 µg/ml) for an hour at 37 ^°^C before adding to the cells.

### Cell lines and cell culture

The cell lines were plated onto cell culture flasks or coverslips, coated with poly-L-lysine, and cultured in a 5 % CO_2_ incubator at 37 ^°^C. The tsA-201 cells (European Collection of Cell Cultures, female sex) were cultured in Dulbecco’s modified Eagle’s medium (DMEM) supplemented with 10 % foetal bovine serum (FBS), 1 unit/ml penicillin, 1 μg/ml streptomycin and 1 % GlutaMAX (Life Technologies, Waltham, MA). When protease inhibitors were used, they were applied 12 h after transfection by replacing the medium with serum-free DMEM F12 (supplemented with 1 unit/ml penicillin, 1 μg/ml streptomycin and 1 % GlutaMAX) containing the inhibitors, as indicated. The cells were incubated in culture for 24 h before harvesting. The production and verification of the CRISPR WT and knockout HEK293 ADAM17^-/-^ and ADAM10^-/-^ cells is described previously (Riethmueller et al., 2016). The SH-SY5Y human neuroblastoma cell line (ECACC # 94030304; female sex) (Biedler et al., 1973) stably expressing HA-α_2_δ-3 was generated in the laboratory by standard techniques, described previously (Davies et al., 2006).

### Cell line transfection

For electrophysiological studies, CRISPR WT and knockout HEK293 cells were transfected with Ca_V_2.2-HA or Ca_V_1.2 together with α_2_δ-1 and β1b or β3 (all in vector pcDNA3) in a ratio 3:2:2. The transfection reagent used was PolyJet (Tebu-bio Ltd), used in a ratio of 3:1 to DNA mix. Culture medium was changed 12 h after transfection and cells were incubated at 37 ^°^C for a further 42 h. CD8 was used as transfection marker.

For cell surface biotinylation and other biochemical experiments, tsA-201 cells were transfected using Fugene6 (Promega) according to the manufacturer’s protocol. CRISPR WT and knockout HEK293cells were transfected with PolyJet as above, and incubated at 37 ^°^C for 48 h.

### Preparation of WCL, deglycosylation, cell surface biotinylation, and immunoblotting

Cell surface biotinylation experiments were carried out on tsA-201 or HEK293 CRISPR WT and knockout cells expressing the cDNAs described. At 48 h after transfection, cells were rinsed with phosphate-buffered saline (PBS) and then incubated for 30 min at room temperature (RT) with 0.5 mg/ml Premium Grade EZ-link Sulfo-NHS-LC-Biotin (Thermo Scientific) in PBS. The reaction was quenched by removing the biotin solution and replacing with PBS containing 200 mM glycine for 2 min at RT. The cells were rinsed with PBS before being resuspended in PBS containing 1 % Igepal; ‘0.1 % SDS and protease inhibitors (PI, cOmplete, Sigma-Aldrich), pH 7.4, for 30 min on ice to allow cell lysis. WCL were then cleared by centrifugation at 13,000 × g and assayed for total protein (Bradford assay, Biorad). Biotinylated lysates were equalised to between 0.5 and 1 mg/ml total protein concentration; 0.5 mg of these biotinylated lysates were adjusted to 500 μl and applied to 40 µl prewashed streptavidin-agarose beads (Thermo Scientific) and rotated overnight at 4 ^°^C. The streptavidin beads were then washed 3 times with PBS containing 0.1 % Igepal, and re-suspended in Peptide N-glycosidase (PNG)-ase-F buffer (PBS, pH 7.4, supplemented with 75 mM β-mercaptoethanol, 1% Igepal, 0.1% SDS, and PI) and de-glycosylated for 3 h at 37 ^°^C with 1 unit of PNGase-F (Roche Applied Science) added per 10 μl volume. When Endo-H was used, a sample of washed beads was removed (before PNGase-F was added), denatured at 99 ^°^C for 10 minutes and treated with Endo-H (New England Biosciences) for 1 h at 37^°^C. Samples were then resuspended in an equal volume of 2 x Laemmli buffer (Davies et al., 2010), supplemented with dithiothreitol (DTT) to a final concentration of 100 mM, and heated for 10 min at 60 ^°^C to elute the precipitated protein. Aliquots of cleared WCL, corresponding to 20 - 40 µg total protein were de-glycosylated in parallel, as described above (Kadurin et al., 2016). The samples were then resuspended in an equal volume of 2 × Laemmli buffer (with 100 mM DTT), followed by 10 min incubation at 60^°^C before loading on SDS-polyacrylamide gel electrophoresis (PAGE). The samples were then resolved by SDS-PAGE on 3 – 8 % Tris-Acetate gels (Thermo Fisher Scientific) and transferred to polyvinylidene fluoride (PVDF) membranes (Biorad). The membranes were blocked with 5 % bovine serum albumin (BSA), 0.5 % Igepal in Tris-buffered saline (TBS) for 30 min at RT and then incubated overnight at 4 ^°^C with the relevant primary Ab. After washing in TBS containing 0.5 % Igepal, membranes were incubated with the appropriate secondary Ab for 1 h at RT. The signal was obtained by HRP reaction with fluorescent product (ECL 2; Thermo Scientific) and membranes were scanned on a Typhoon 9410 phosphorimager (GE Healthcare).

### Preparation of Whole Cell Membrane fraction and of Detergent Resistant Membranes (DRMs)

Cell pellets of a confluent T-75 flask were resuspended in 1 ml of ice-cold buffer containing 10 mM NaCl, 10 mM HEPES, pH 7.4, and PI. Cells were lysed by 10 passages through a 25-gauge syringe, followed by three 10 s rounds of sonication. Cell debris was removed by centrifugation (1000 × g, for 10 min at 4 ^°^C), and the resultant supernatants were recentrifuged (60,000 × g, for 60 min at 4 ^°^C) to pellet membranes. The whole cell membrane fraction was resuspended in 50 µl of PNGase-F buffer and de-glycosylated for 3 h at 37 ^°^C with 1 unit of PNGase-F added per 10 μl volume.

The protocol for preparation of DRMs was similar to that described previously (Davies et al., 2010; Kadurin et al., 2012). All steps were performed on ice. Confluent tsA-201 cells from two 175 cm^2^ flasks were taken up in Mes-buffered saline (MBS, 25 mm Mes, pH 6.5, 150 mm NaCl, and PI) containing 1% (v/v) Triton X-100 (TX-100) (Thermo Scientific), and left on ice for 1 h. An equal volume of 90% (w/v) sucrose in MBS was then added to a final concentration of 45 % sucrose. The sample was transferred to a 13 ml ultracentrifuge tube and overlaid with 10 ml of discontinuous sucrose gradient, consisting of 35% (w/v) sucrose in MBS (5 ml) and 5% (w/v) sucrose in MBS (5 ml). The sucrose gradients were ultracentrifuged at 140,000 x *g*_avg_ (Beckman SW40 rotor) for 18 h at 4 ^°^C. 1 ml fractions were subsequently harvested from the top to the bottom of the tube and aliquots of 10 μl from each fraction were analysed by SDS-PAGE and western blotting to obtain DRM profiles. When necessary, DRMs (combined peak fractions identified by presence of flotillin-1) from the gradient were washed free of sucrose by dilution into 25 volumes of cold PBS (pH 7.4) and pelleted by ultracentrifugation at 150,000 × *g* (Beckman Ti 70 rotor) for 1 h at 4 ^°^C. TX-100-insoluble protein was resuspended in PNGase-F buffer and de-glycosylated for 3 hr at 37^°^C with 1 unit of PNGase-F added per 10 μl volume. The samples were then resuspended in Laemmli buffer (1 x final concentration, with 100 mM DTT) followed by 10 min incubation at 60 ^°^C before loading on SDS-PAGE.

### Subcellular fractionation of SH-SY5Y cell line stably expressing HA-α_2_δ-3 subunits

The subcellular fractionation was performed in a continuous iodixanol gradient within the range 0-25% (w/v) iodixanol to resolve the major membrane compartments of the ER, Golgi membranes from a post nuclear supernatant prepared from a cultured cell homogenate, as described previously (Graham, 2002). Briefly, two T-175 flasks of SH-SY5Y cells (∼ 70 % confluent) stably expressing HA-α_2_δ-3 were washed with PBS, and suspended in 3 ml of Homogenisation Medium (HM, 0.25 M sucrose, 1 mM EDTA 10 mM Tris, pH 7.4; supplemented with PI). Cell pellets were disrupted using a 25-gauge syringe (5 x passes), and centrifuged at 1,000 x g for 10 min. A 25% (w/v) iodixanol solution was prepared by mixing equal volumes of HM and Working Solution (5 vol of OptiPrep™ + 1 vol of diluent (0.25 M sucrose, 6 mM EDTA, 60 mM Tris, pH 7.4; supplemented with PI). A 10 ml gradient was prepared in Beckman SW40 rotor tubes, using equal volumes of HM and the 25% iodixanol solution using a two-chamber gradient maker. 1 ml of the supernatants from the 1000 x g centrifugation were laid on top of the gradient and centrifuged at 200,000 x g (Beckman SW40 rotor) for 4 h. Fractions (0.75 ml) were collected from the top. Aliquots of each fraction were supplemented with Triton X100 to 0.5 %, SDS to 0.1 %, β-mercaptoethanol to 75 mM and de-glycosylated with PNGase-F as described above. 5 x Laemmli buffer was then added (1 x final concentration, with 100 mM DTT) followed by 10 min incubation at 60 ^°^C before loading on SDS-PAGE.

### Electrophysiology

Calcium channel currents in transfected HEK293 CRISPR WT and knockout cells were investigated by whole cell patch-clamp recording, essentially as described previously (Berrow et al., 1997). The patch pipette solution contained in mM: Cs-aspartate, 140; EGTA, 5; MgCl_2_, 2; CaCl_2_, 0.1; K_2_ATP, 2; Hepes, 20; pH 7.2, 310 mOsm with sucrose. The external solution for recording Ba^2+^ currents contained in mM: tetraethylammonium (TEA) Br, 160; KCl, 3; NaHCO_3_, 1; MgCl_2_, 1; Hepes, 10; glucose, 4; BaCl_2_, (1 for Ca_V_2.2 currents and 5 for Ca_V_1.2 currents); pH 7.4, 320 mosM with sucrose. An Axopatch 1D or Axon 200B amplifier was used, and whole cell voltage-clamp recordings were sampled at 10 kHz frequency, filtered at 2 kHz and digitized at 1 kHz. 70-80 % series resistance compensation was applied and all recorded currents were leak subtracted using P/8 protocol. Membrane potential was held at −80 mV. Analysis was performed using Pclamp 9 (Molecular Devices) and Origin 2017 (Microcal Origin, Northampton, MA). Current-voltage (*I-V*) relationships were fit by a modified Boltzmann equation as follows: *I = G*_*max*_*\*(V-V*_*rev*_*)/(1+exp(-(V-V*_*50, act*_*)/k))*, where *I* is the current density (in pA/pF), *G*_max_ is the maximum conductance (in nS/pF), *V*_rev_ is the apparent reversal potential, *V*_50, act_ is the midpoint voltage for current activation, and *k* is the slope factor.

### Immunocytochemistry, imaging and analysis

Immunocytochemistry was carried out on HEK293 CRISPR WT and knockout cells expressing GFP_Ca_V_2.2-HA together with α_2_δ-1 and β1b. After transfection, cells were incubated for 48 h before being fixed with 4 % paraformaldehyde (PFA) in PBS, pH 7.4 at RT for 5 min. Blocking was performed for 30 min at RT in PBS containing 20 % goat serum and 5 % BSA. An anti-HA Ab (rat monoclonal) was applied (100 ng/ml dilution in PBS with 10 % goat serum and 2.5 % BSA) for 1 h at RT to the non-permeabilized cells. Cells were then incubated with an anti-rat Alexa Fluor 594 (1:500 dilution in PBS, containing 2.5 % BSA and 10 % goat serum) at RT for 1 h. The coverslips were mounted onto glass slides using VECTASHIELD® mounting medium (Vector Laboratories, Peterborough, UK).

Imaging was performed on Zeiss LSM 780 confocal microscope, at fixed microscope settings for all experimental conditions of each experiment. Images of HEK293 CRISPR WT and knockout cells were obtained using a 63 x oil objective at a resolution of 1,024 × 1,024 pixels and an optical section of 0.5 μm. After choosing a region of interest containing transfected cells, the 3 x 3 tile function of the microscope allowed imaging of a larger area selected without bias. Every cell identified as transfected was included in the measurements, to ensure lack of bias.

Images of HEK293 CRISPR WT and knockout cells were analyzed using ImageJ (*imagej*.*net*). Cell surface signal was quantified using the freehand line (3 pixels) to trace the membrane region stained by anti-HA Ab. The total level of Ca_V_2.2 corresponding to the GFP signal was measured using the freehand selection tool, excluding the nucleus. The value of the mean pixel intensity in different channels was measured separately and background was subtracted by measuring the intensity of an imaged area without transfected cells. The ratio of cell surface to total Ca_V_2.2 (HA/GFP) was then calculated for each cell. The data are shown as mean ± SEM and single data points (for 6 independent transfections, in which all conditions were examined in parallel).

### Quantification and statistical analysis

Data were analyzed with GraphPad Prism 8 (GraphPad software, San Diego, CA) or Origin-Pro 2017 (OriginLab Corporation, Northampton, MA, USA). All data are shown as mean ± SEM; “n” refers to number of experiments, unless indicated otherwise, and is given in the figure legends, together with details of statistical tests used. Experiments where representative data are shown were repeated at least 3 times, as stated. Graphpad Prism 8 was used for statistical analysis. Statistical significance between two groups was assessed by Student’s t test, as stated. One-way or two-way ANOVA and the stated post-hoc analysis was used for comparison of means between three or more groups.

## ACKNOWLEDGEMENTS

We thank Prof. Paul Saftig (Biochemisches Institut, Christian-Albrechts-Universität Kiel, Kiel, Germany) for the generous gift of the ADAM17 and ADAM10 knockout and CRISPR WT HEK293 cells, and for very useful advice during the course of this project.

The work of the ACD was supported by a Wellcome Trust Investigator award 206279/Z/17/Z, and the work of IK was supported in part by British Heart Foundation grant PG/18/83/34123.

## AUTHOR CONTRIBUTIONS

IK performed all biochemical experiments except those in Figure 5, performed by KMP. SD performed all electrophysiology (Figure 3), and imaging (Figure 4). KMP made cDNA constructs. KC performed tissue culture of cell lines and made stable SH-SY5Y cell line. JIJE performed initial experiments relating to Figure 2. LT and HN provided reagents and ideas for experiments, and reviewed data. ACD and IK conceived the study and ACD, IK, KMP and SD wrote the manuscript aided by all the other authors.

## SUPPLEMENTAL INFORMATION

### Supplemental Figures

**Figure S1 (relates to Figure 2).**
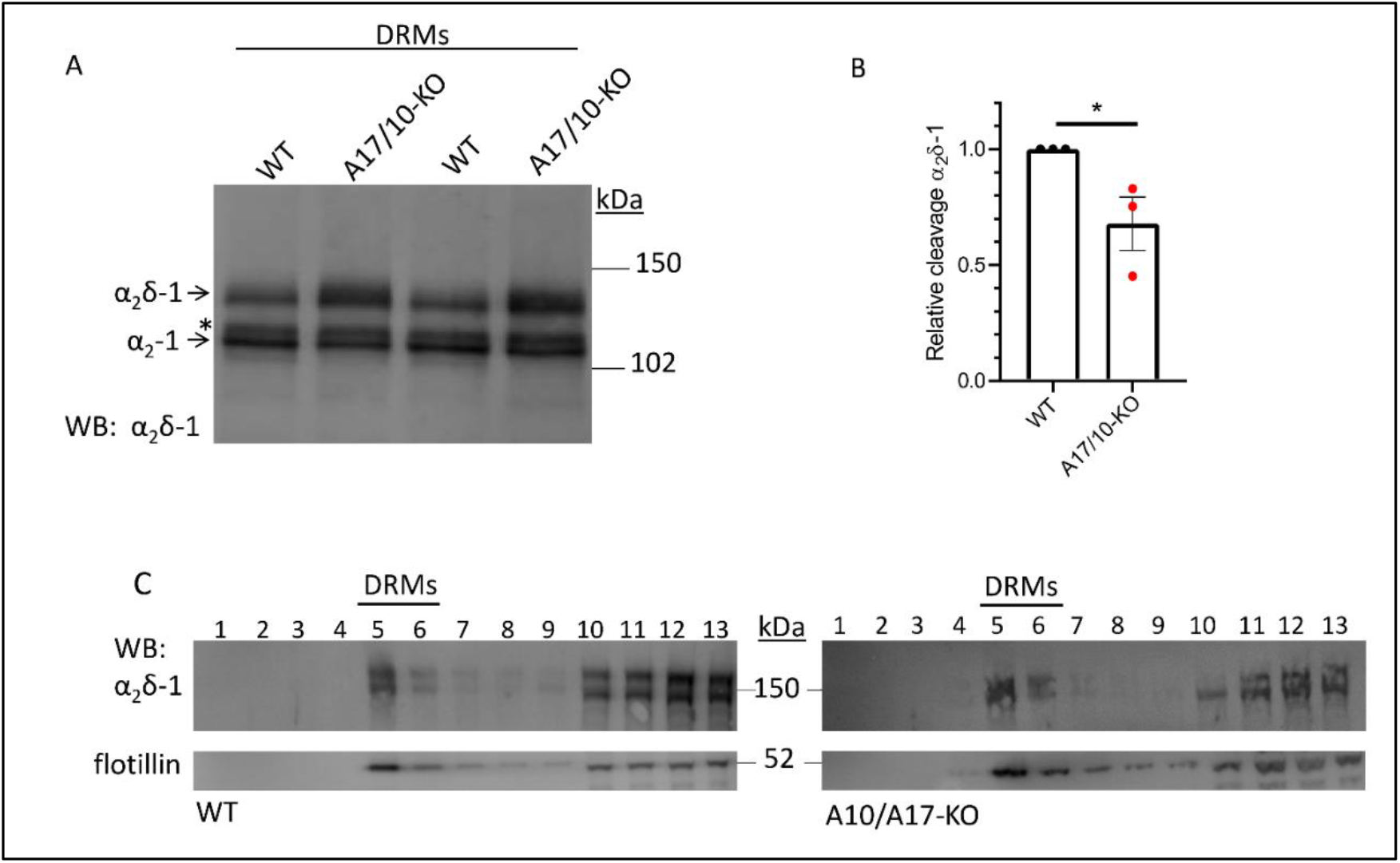
Cleavage of α_2_δ-1 is reduced in DRMs from CRISPR *ADAM17*^-/-^ / *ADAM10*^-/-^ compared to CRISPR WT cells. (A) Effect of expression of α_2_δ-1 in CRISPR WT (lanes 1 and 3) compared to *ADAM17*^-/-^ / *ADAM10*^-/-^ (A17/10-KO) HEK293 cells (lanes 2 and 4) on cleavage of HA-α_2_δ-1 in DRM fraction (α_2_-1 immunoblot), de-glycosylated to allow resolution between pro-α_2_δ-1 (upper band) and the cleaved form, α_2_-1 (lower band). * indicates an intermediate species, which may represent cleavage of α_2_δ-1 at an alternative site, or an intermediate product. (B) Quantification of the effect of expression in *ADAM17*^-/-^ / *ADAM10*^-/-^ cells on relative cleavage of α_2_δ-1 in DRMs (normalized to that under control conditions). The data are mean ± SEM and individual data from 3 separate experiments. Statistical difference determined using Student’s t test; * *P* = 0.049. In this study there was ∼70% absolute cleavage of α_2_δ-1 in DRMs from CRISPR WT cells and ∼45% cleavage in *ADAM17*^-/-^ / *ADAM10*^-/-^ cells. (C) Sucrose gradient profiles (lanes 1-13) showing α_2_δ-1 distribution (upper panels) from CRISPR WT (left) compared to *ADAM17*^-/-^ / *ADAM10*^-/-^ HEK293 cells (right). Peak DRM fractions (5 and 6) shown by bar, and by presence of flotillin (lower panels).

**Figure S2 (relates to Figure 5).**
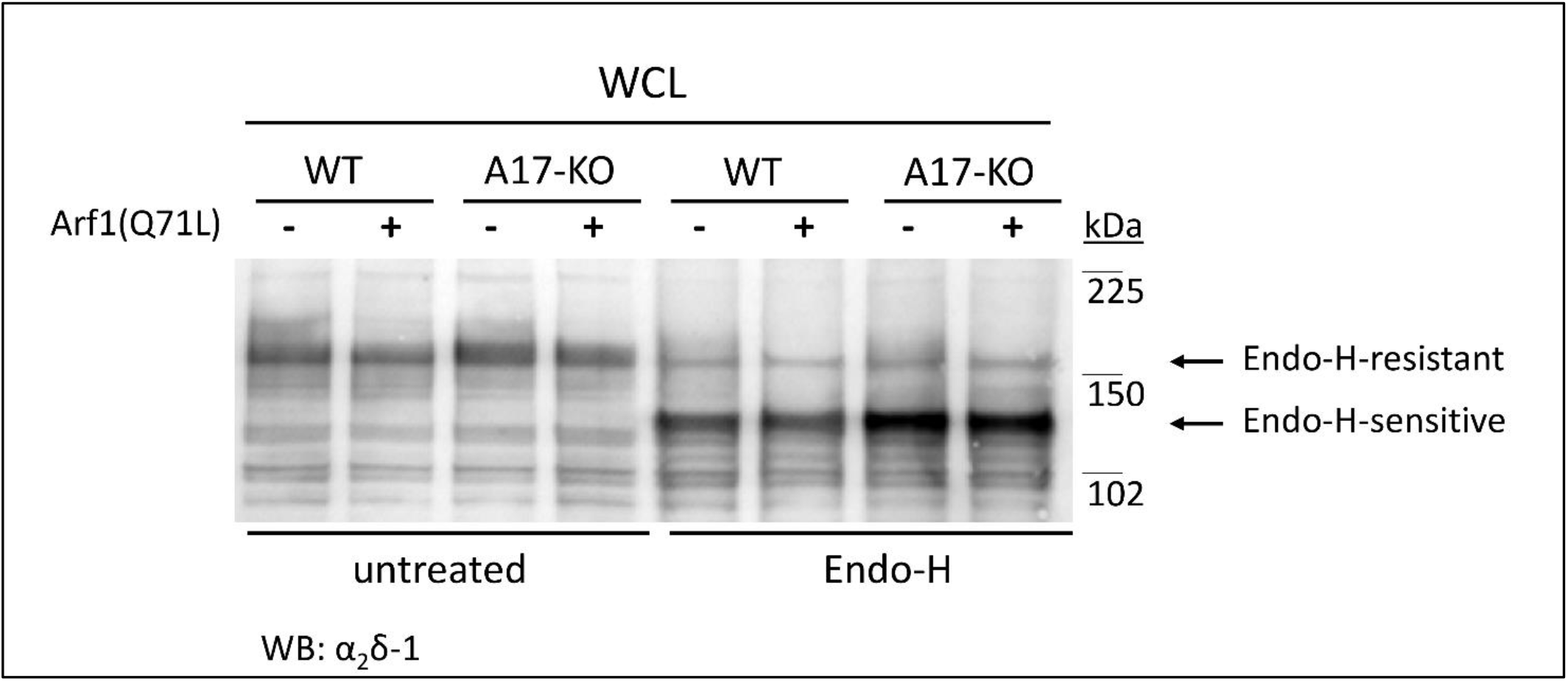
Most α_2_δ-1 in WCL is Endo-H-sensitive in both CRISPR WT and *ADAM17*^-/-^ cells. Effect of expression of α_2_δ-1 in CRISPR WT (lanes 1, 2, 5, 6) and *ADAM17*^-/-^ (A17-KO) cells (lanes 3, 4, 7, 8) in the absence (lanes 1, 3, 5, 7) and presence (lanes 2, 4, 6, 8) of the ER-to-Golgi blocker, Arf1(Q71L), in WCL samples either left untreated (lanes 1-4) or treated with Endo-H (lanes 5-8) for comparison. The sizes of the Endo-H-resistant bands and Endo-H-sensitive bands are indicated with arrows. Most α_2_δ-1 in the WCL is Endo-H-sensitive (lower band), suggesting that it is in the ER and has not yet progressed through the Golgi complex.

**Figure S3 (relates to Figure 5).**
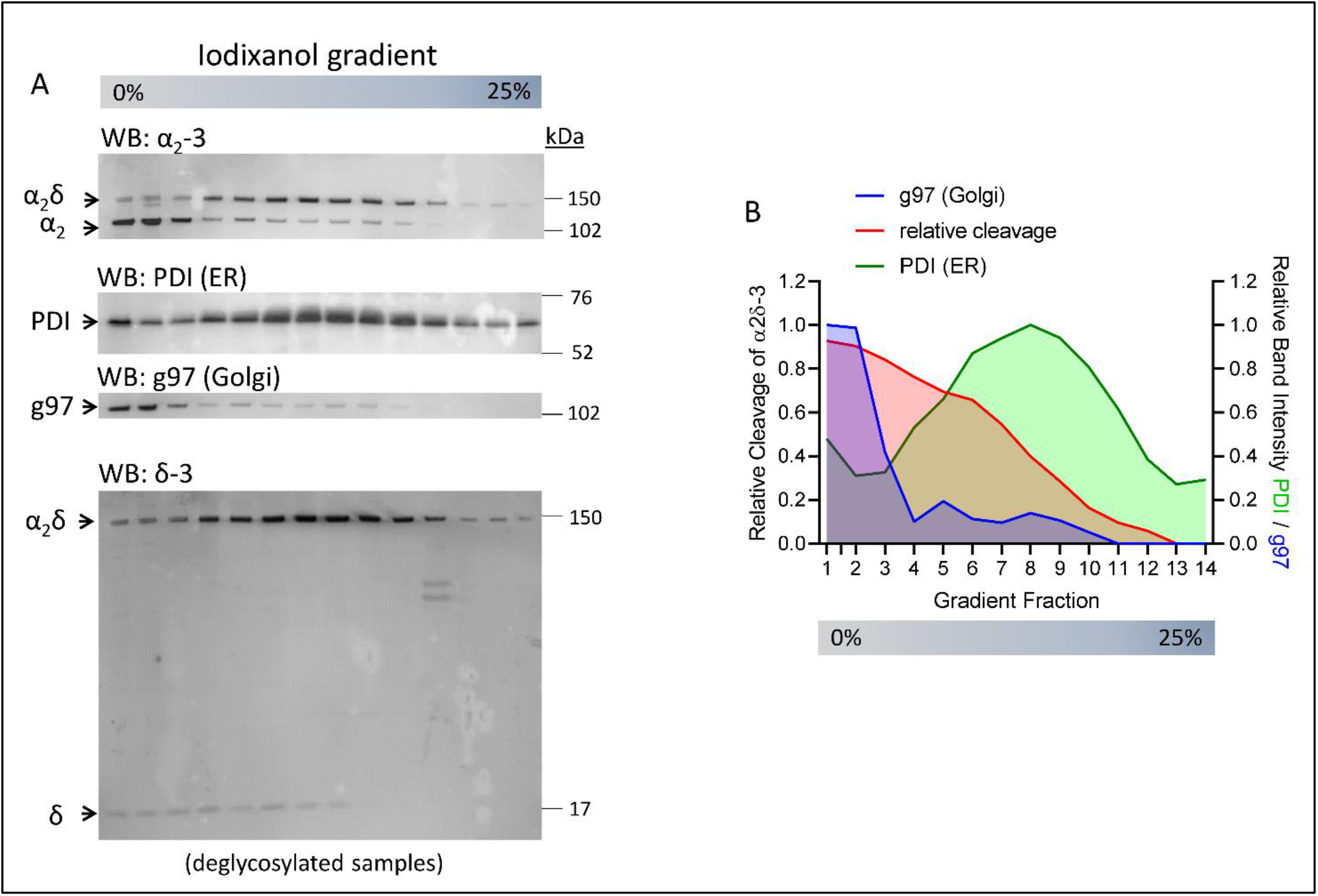
Gradient centrifugation of α_2_δ-3 stable cell line shows proteolytic cleavage appears associated with the Golgi fraction. (A) Distribution of α_2_δ-3 (α_2_-3 Ab, top panel, showing uncleaved α_2_δ-3 and cleaved α_2_-3, as indicated) following iodixanol gradient centrifugation of SH-SY5Y cells, compared to the ER marker Protein disulfide isomerase (PDI, second panel) and the Golgi marker Golgin 97 (g97, third panel) and δ-3 (bottom panel, showing uncleaved α_2_δ-3 and cleaved δ-3, as indicated). (B) Plot of relative cleavage of α_2_δ-3 (red) compared to distribution of g97 (purple) and PDI (green) along iodixanol gradient.

